# PME-1 suppresses anoikis and is associated with therapy relapse of PTEN-deficient prostate cancers

**DOI:** 10.1101/581660

**Authors:** Anna Aakula, Aleksi Isomursu, Christian Rupp, Andrew Erickson, Otto Kauko, Pragya Shah, Artur Padzik, Yuba Raj Pokharel, Amanpreet Kaur, Song-Ping Li, Lloyd Trotman, Pekka Taimen, Antti Rannikko, Jan Lammerding, Ilkka Paatero, Tuomas Mirtti, Johanna Ivaska, Jukka Westermarck

**Affiliations:** Turku Bioscience Centre, University of Turku and Åbo Akademi University, Turku, Finland; HUSLAB Laboratory Services, Helsinki University Hospital Medicum and Institute for Molecular Medicine Finland FIMM, University of Helsinki, Helsinki, Finland; Weill Institute for Cell and Molecular Biology & Meinig School of Biomedical Engineering, Cornell University, Ithaca, NY, USA; Institute of Biomedicine, University of Turku, Turku, Finland; Cold Spring Harbor Laboratory, Cold Spring Harbor, NY, USA; Department of Pathology, Turku University Hospital, Turku, Finland; Department of Urology, Helsinki University Central Hospital, Helsinki, Finland; Department of Biochemistry, University of Turku, Turku, Finland; InFLAMES Research Flagship Center, University of Turku, Turku, Finland

**Keywords:** AR, ERG, integrin, extracellular matrix (ECM), nuclear lamina, LAP2

## Abstract

While organ-confined PCa is mostly therapeutically manageable, metastatic progression of PCa remains an unmet clinical challenge. Resistance to anoikis, a form of cell death initiated by cell detachment from the surrounding extracellular matrix, is one of the cellular processes critical for PCa progression towards aggressive disease. Therefore, further understanding of anoikis regulation in PCa might provide therapeutic opportunities. Here, we discover that PCa tumors with concomitantly compromised function of two tumor suppressor phosphatases, PP2A and PTEN, are particularly aggressive, having less than 50% 5-year secondary-therapy free patient survival. Functionally, overexpression of PME-1, a PP2A inhibitor protein, inhibits anoikis in PTEN-deficient PCa cells. *In vivo,* PME-1 inhibition increased apoptosis in *in ovo* PCa tumor xenografts, and attenuated PCa cell survival in zebrafish circulation. Molecularly, PME-1 deficient PCa cells display increased trimethylation at lysines 9 and 27 of histone H3 (H3K9me3 and H3K27me3), a phenotype corresponding to increased apoptosis sensitivity. In summary, we discover that PME-1 overexpression supports anoikis resistance in PTEN-deficient PCa cells. Clinically, the results identify PME-1 as a candidate biomarker for a subset of particularly aggressive PTEN-deficient PCa.

## INTRODUCTION

Prostate cancer (PCa) is often detected early, and can remain non-aggressive, and non-metastatic for years [1, 2]. Whereas local PCa is not life-threatening, transformation from indolent to aggressive disease is clinically the most important transition in PCa progression. Thereby understanding this switch in more detail might provide therapeutic opportunities. Analysis of the evolutionary history of lethal metastatic PCa has revealed that tumor suppressor proteins are particularly lost during transition towards metastatic human PCa [1]. One such tumor suppressor is a dual-specificity phosphatase Phosphatase and Tensin homolog (PTEN) that is inactivated in a large fraction of high grade PCas [3]. Both in human and mice, prostate-specific PTEN deletion leads to an aggressive and metastatic PCa [3–5]. Protein phosphatase 2A (PP2A) is another tumor suppressor phosphatase that is commonly inactivated in aggressive PCa. Specifically, PP2A B-subunits PPP2R2A and PPP2R2C (B55α and B55γ) have been reported as either genetically deleted, or down-regulated in human PCa samples [6–8]. PP2A is inactivated also non-genetically in human cancers by overexpression of inhibitor proteins such as CIP2A, PME-1 or SET [9, 10]. CIP2A overexpression clinically associates with castration-resistant prostate cancer (CRPC) [11], and inhibition of both CIP2A and SET inhibits malignant growth of PCa cells [11, 12]. In contrast, the role of PME-1 in PCa is currently unknown. PME-1 inhibits PP2A activity both by directly binding to catalytic centre of the catalytic subunit PP2A-C, and by inhibiting recruitment of tumor suppressive B-subunits to the PP2A complex [10, 13]. Thereby, PME-1 expression levels provide an indirect read-out for effective PP2A inhibition in cells and in tissues.

One of the hallmarks of PCa progression towards aggressive CRPC is acquisition of resistance towards a specific type of programmed cell death, anoikis [14]. Anoikis is induced by detachment of cells from other cells, or the surrounding extracellular matrix (ECM). Anoikis suppression is not only relevant for indolent PCa cells to acquire anchorage-independence, but also for survival of PCa cells with metastatic potential in circulation [15, 16]. Thus, understanding the mechanisms driving anoikis resistance of PCa cells could provide novel therapy opportunities for clinical management of PCa by facilitating inhibition of progression to metastatic disease. Anoikis resistance in PCa has been linked to changes in cell adhesion, cytoskeleton, epigenetics, as well as deregulated intracellular survival pathways [14]. Activated integrin signaling is a central contributor to anchorage-independence and metastasis [17, 18], and integrin activation correlates with anoikis resistance in PCa cell lines [19, 20]. Recently, epigenetic regulation of histone H3 methylation has also surfaced as an important mechanism regulating apoptosis and anoikis [21–23]. The role of PP2A in PCa anoikis sensitivity is, however, poorly understood.

In this study, we demonstrate that PP2A inhibitor protein PME-1, that has not been previously implicated in PCa, has a critical role in anoikis suppression of PTEN-deficient PCa cells both *in vitro* and *in vivo.* Clinically, PCa tumors with inhibition of both tumor suppressor phosphatases PTEN (genetic deletion) and PP2A (PME-1 overexpression) had remarkably short relapse-free survival.

## RESULTS

### PME-1 overexpression in prostate cancer associates with PTEN loss and with therapy relapse

PME-1 protein expression and its clinicopathological associations were evaluated in PCa tissue microarray (TMA) material consisting of 358 patients treated primarily with radical prostatectomy in the Helsinki University Hospital between 1983 and 1998. The clinical cohort has previously been described in detail [5, 24] (see Table S1 for demographics). The specificity of the PME-1 antibody for immunohistochemical (IHC) staining has been validated previously [25]. PME-1 expression was scored using a 4-tier scale of negative, low, intermediate, and strong expression. As each patient had three cancerous cores in the TMA, the maximum values of PME-1 scores were used and each patient was dichotomized as either high or low (negative vs. intermediate to strong staining) (Figure 1A). In the correlation analysis with clinical variables, strong PME-1 protein expression correlated with higher grade group and advanced stage (Table 1). We also correlated PME-1 status with previously assessed PTEN, ERG and AR status [5, 24]. Notably, high PME-1 expression significantly associated with complete PTEN loss, but also with ERG positivity and high AR expression status (Figure S1A).

**Figure 1.**
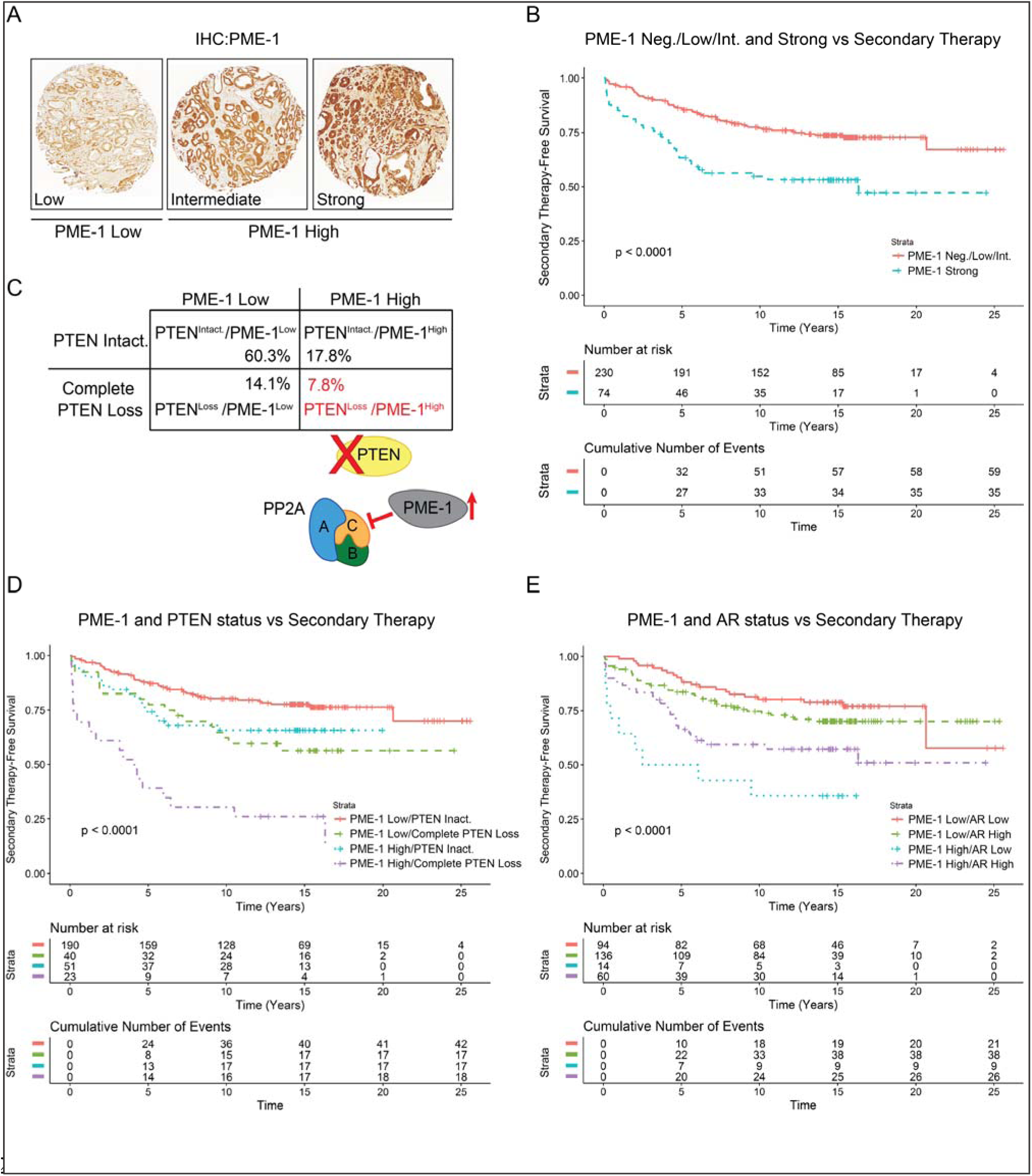
PME-1 overexpression associates with PTEN loss and therapy relapse of PTEN negative prostate cancer patients. **A**. PME-1 protein expression in PCa tissue microarray material from 358 patients treated primarily with radical prostatectomy in the Helsinki University Hospital between 1983 and 1998 was assessed by immunohistochemical (IHC) stainings. PME-1 protein content was scored using 4-tier scale; negative (not shown), low, intermediate and strong expression. **B**. Kaplan-Meyer analysis of time to secondary therapies after primary treatment based on PME-1 status alone. **C**. Assessment of the status of PME-1 and PTEN in the patient populations and the putative effect on PP2A activity**. D-E**. Kaplan-Meyer analysis of time to secondary therapies after primary treatment based on PME-1 status in combination with PTEN deletion (D) and AR expression (E).

Patients with high tumor PME-1 expression had significantly shorter time to secondary therapies after primary treatment (i.e., relapse-free survival) (Figure 1B), indicative of clinical relevance of PME-1 in human PCa. Linked to the association between PTEN loss and high PME-1 expression (Figure S1A), we asked whether patients with such tumors, having both tumor suppressor phosphatases (PTEN and PP2A) inhibited, would present with a particularly aggressive disease? Patients with PTEN^loss^/PME-1^high^ tumor phenotype constituted a sub-cohort of approximately 8% of patients (Figure 1C). Strikingly, PTEN^loss^/PME-1^high^ status correlated with remarkably aggressive tumors, with about 40% 5-year secondary-therapy free survival among the cohort (Figure 1D). Similar, albeit less prominent co-operative effects were observed with PME-1 overexpression and with either AR (Figure 1E) or ERG expression (Figure S1B).

These results reveal a clinical relevance of PME-1 in PCa. Notably, they identify a very aggressive PCa subpopulation associated with concomitant inhibition of PTEN and PP2A tumor suppressor phosphatases.

### PME-1 promotes anchorage-independent growth of prostate cancer cells

Acknowledging the clinical indications for a link between PME-1 and PTEN, we tested the impact of PME-1 depletion on colony growth of two PTEN-deficient PCa cells lines, PC-3 (PTEN null) and DU-145 (PTEN heterozygous). PME-1 silencing by two independent siRNAs did not affect colony growth of either of the cell lines in 2D adherent cell culture conditions (Figure 2A). However, PME-1 silencing decreased anchorage-independent growth of both cell lines in soft agar (Figure 2B). The context-dependent role of PME-1 in supporting anchorage-independent growth, but being redundant for cell growth in 2D adherent conditions is fully consistent with the established requirement of PP2A inhibition for anchorage-independence of transformed human cells [26].

**Figure 2.**
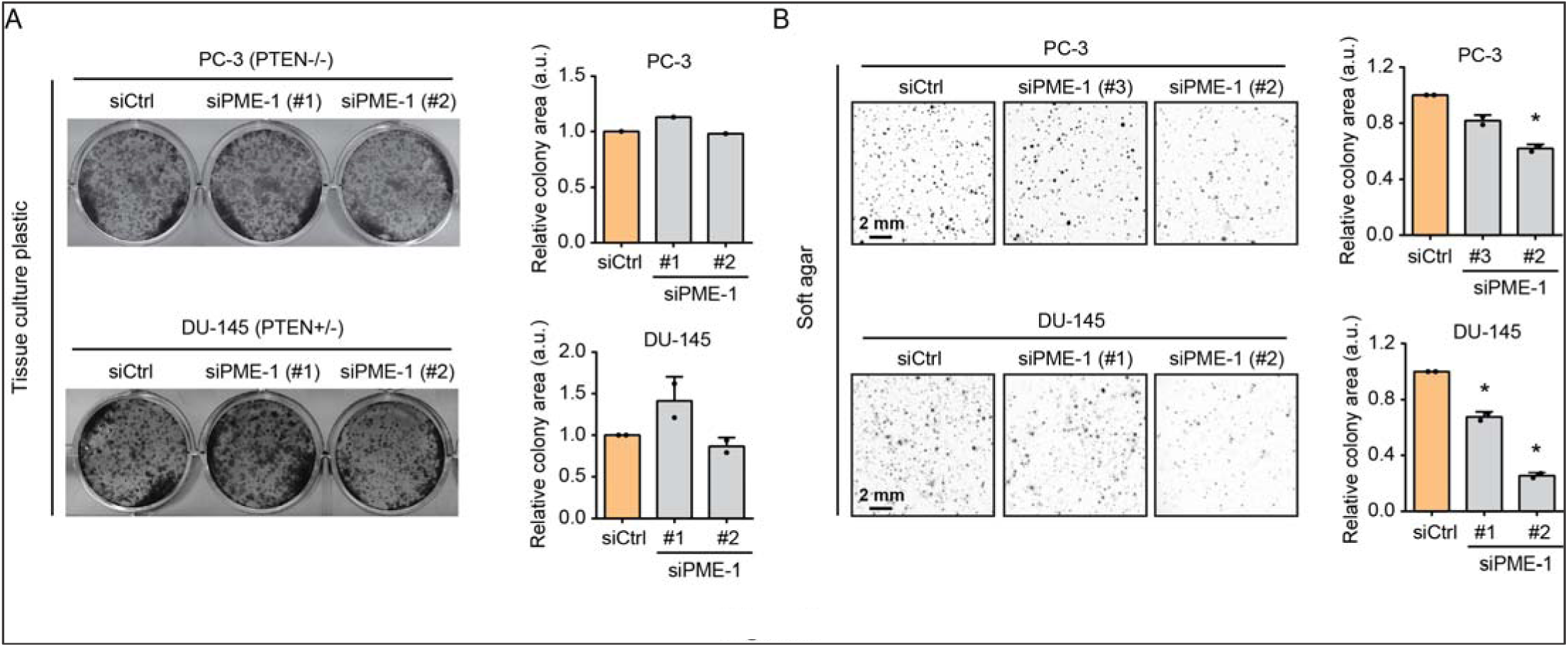
PME-1 promotes anchorage-independent growth of prostate cancer cells. **A**. The effect of PME-1 depletion, utilizing two independent siRNAs, was investigated by colony formation assays (10 days of growth) in two PTEN-deficient PCa cells lines PC-3 and DU-145. (Left) Representative images of the wells. (Right) Bar graphs depicting the quantified data, mean ± SD of one to two independent experiments. **B**. The effect of PME-1 knock-down on anchorage-independent growth in soft agar assays (14 days of growth) in both PC-3 and DU-145 cells. (Left) Representative images depicting the colonies. (Right) Bar graphs displaying the quantified data, mean ± SD of two independent experiments. *p < 0.05, Welch’s t-test.

To further assess whether PME-1 is particularly relevant for survival under anchorage-independent conditions, we evaluated apoptosis induction in control and PME-1 siRNA transfected PC-3 cells cultured either on tissue culture plastic or low attachment plates. We observed a strong synergy between PME-1 inhibition (by two independent siRNA oligos) and low attachment culture conditions in apoptosis induction measured by PARP cleavage (Figure 3A). To further rule out the possibility of siRNA off-target effects, we created PC-3 PME-1 knock-out (KO) cells (pool or single cell clone) by CRISPR/Cas9. These cells displayed dose-dependent apoptosis induction upon PME-1 loss and following detachment (Figure 3B). The functional co-operation of PTEN and PP2A was interrogated further in a genetically defined model system: PME-1 cDNA overexpression in PTEN/p53 KO mouse embryonic fibroblasts (MEFs) [27]. In accordance with siRNA and CRISPR/Cas9 results, overexpression of PME-1 in PTEN/p53 KO MEFs prevented PARP cleavage upon 24 h incubation in low attachment conditions (Figure 3C). Taken together, these data indicate an important role of PME-1 in supporting anchorage-independent growth of PTEN-deficient PCa cells.

**Figure 3.**
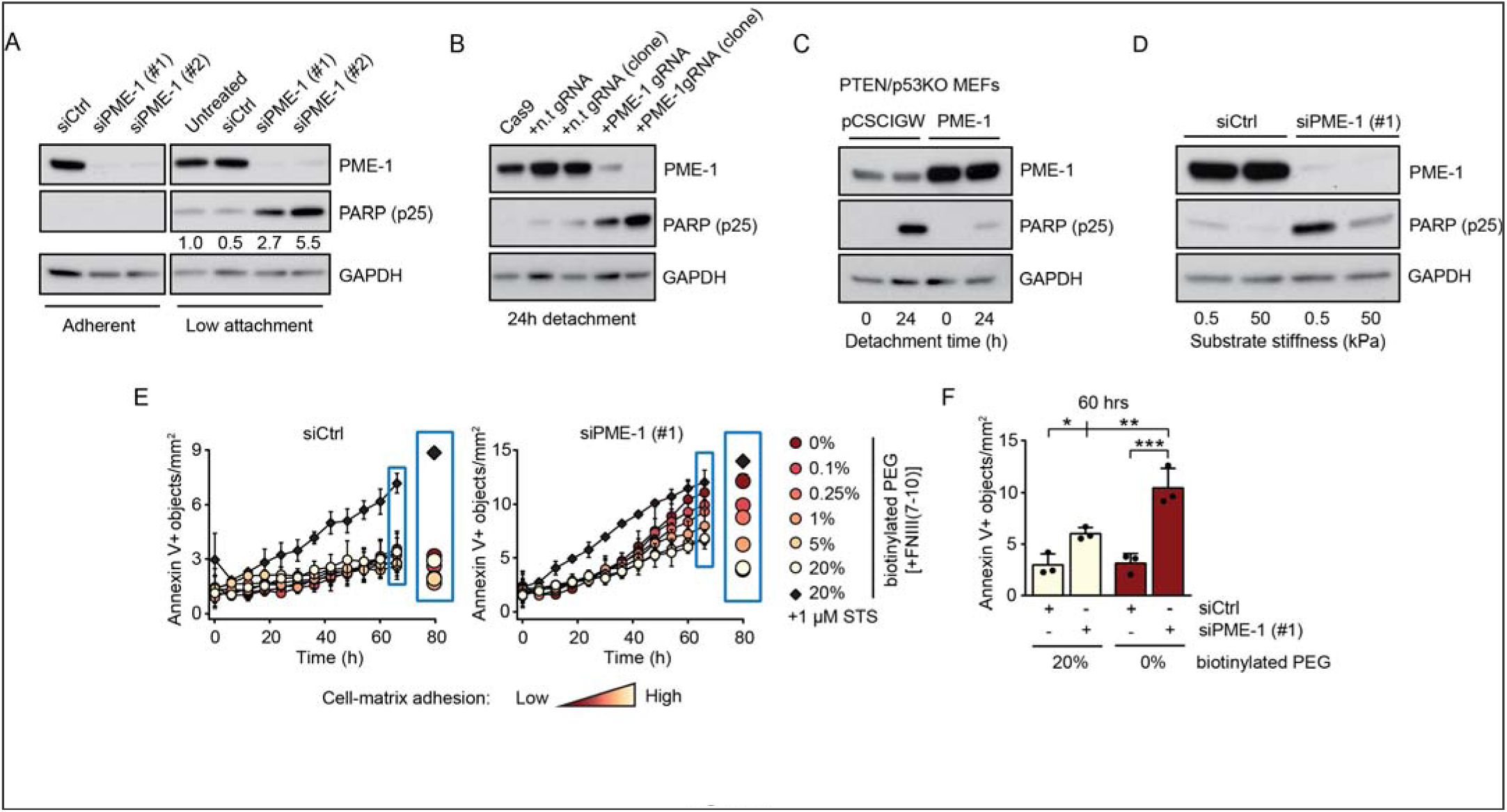
PME-1 promotes anoikis resistance in PTEN-deficient prostate cancer cells. **A**. siCtrl- and siPME-1-transfected PC-3 cells were plated 72 h post-transfection on normal cell culture or low attachment plates for 24 h, before collection and lysis, and subsequently analyzed by western blotting for cleaved PARP-1 (cPARP (p25)). **B**. Apoptosis induction, as measured by PARP cleavage, in CRISPR/Cas9 generated PC-3 cells, with and without non-targeting gRNA (lanes 1-3), a pool of PME-1 gRNA transfected cells (lane 4), and a single cell subclone of PME-1 targeted cells (lane 5), after 24 h detachment. **C.** The effect of stable PME-1 overexpression in mouse embryonic fibroblasts (MEFs) from a PTEN/p53KO mouse model, after 24 h detachment, was assessed by western blotting for cleaved PARP. **D**. Mechanosensitive PARP cleavage in PME-1-depleted PC-3 cells. The cells were cultured on soft (0.5 kPa) or stiff (50 kPa) polyacrylamide hydrogel for 24 h before being scraped into PBS, spun down, lyzed and analyzed for protein expression. **E.** The number of Annexin V positive objects, a marker for apoptosis, in cultures of siCtrl- and siPME-1- transfected PC-3 cells upon varying degrees of cell-matrix interaction. The cells were grown on PEG-coated surfaces with 0-20% biotinylated compound bound to integrin ligand, fibronectin fragment FNIII(7-10). STS, staurosporine. Mean ± SD of three independent experiments**. F.** Annexin V positive objects in PC-3 cultures with no or high levels of integrin ligand, after 60 h of incubation. Mean ± SD of three independent experiments. *p < 0.05, **p < 0.01, ***p < 0.001, One-way ANOVA with Bonferroni’s multiple comparisons test.

In addition to low attachment conditions, culturing cells on low stiffness matrix mimics a situation where cells are loosely associated with their surroundings (i.e., anchorage-independent growth). Previously, it was shown that non-transformed cells succumb to anoikis-type cell death when plated on soft substrates, whereas cancer cells do not die under the same conditions [28, 29]. To test whether PME-1 expression contributes to apoptosis resistance of cancer cells on soft matrix, either control or PME-1 siRNA transfected PC-3 cells were plated on low (0.5 kPa) or high (50 kPa) stiffness hydrogels functionalized with ECM components (fibronectin and collagen I). Consistent with published results from other cancer cells [28], control PC-3 cells showed only a very small increase in PARP cleavage on soft substrate after 24 h. However, PME-1 depletion sensitized the cells to apoptosis induction selectively in low stiffness conditions (Figure 3D). Integrin-mediated cell adhesion to ECM ligands activates outside-in signaling pathways to promote cell proliferation and viability [17, 19]. To test if the apoptosis of PME-1-depleted cells scales with the degree of adhesion signaling, we exposed siCtrl and siPME-1 transfected cells to surfaces functionalized with increasing concentrations of integrin ligand (fibronectin-fragment FNIII(7-10)), while non-specific interactions with the plastic surface were prevented using an anti-fouling agent (Figure S2A). Consequentially, cell adhesion to the surface was fully dependent on the concentration of the integrin ligand (Figure S2B). Apoptosis (Annexin V signal) was highest in PME-1-silenced cells with no available integrin ligand (0%) and decreased proportionally with increasing integrin-mediated adhesion, whereas siCtrl cells were insensitive to lack of adhesion and showed marked apoptosis only upon staurosporine (STS) treatment, which was included as a positive control (Figure 3E,F and S2C).

In summary, these results demonstrate the potential of PME-1 overexpression in protecting PTEN-deficient PCa cells from anoikis.

### PME-1 supports *in vivo* anoikis resistance and survival of prostate cancer cells in circulation

To test the *in vivo* relevance of PME-1-mediated inhibition of PCa cell anoikis, we grew either siCtrl or siPME-1 transfected PC-3 xenografts on chicken embryo chorioallantoic membranes (CAM). Tumors formed by PME-1-depleted cells were overall more translucent, suggesting decreased tumor growth (Figure 4A). In accordance with the anoikis suppressing activity of PME-1, histological analyses of the dissected tumors revealed increased TUNEL positivity in tumors derived from siPME-1 transfected cells (Figure 4B,C). Tumor suppression was further confirmed by the reduced number of Ki-67 positive cells in tumors derived from siPME-1 transfected cells (Figure 4B,C).

**Figure 4.**
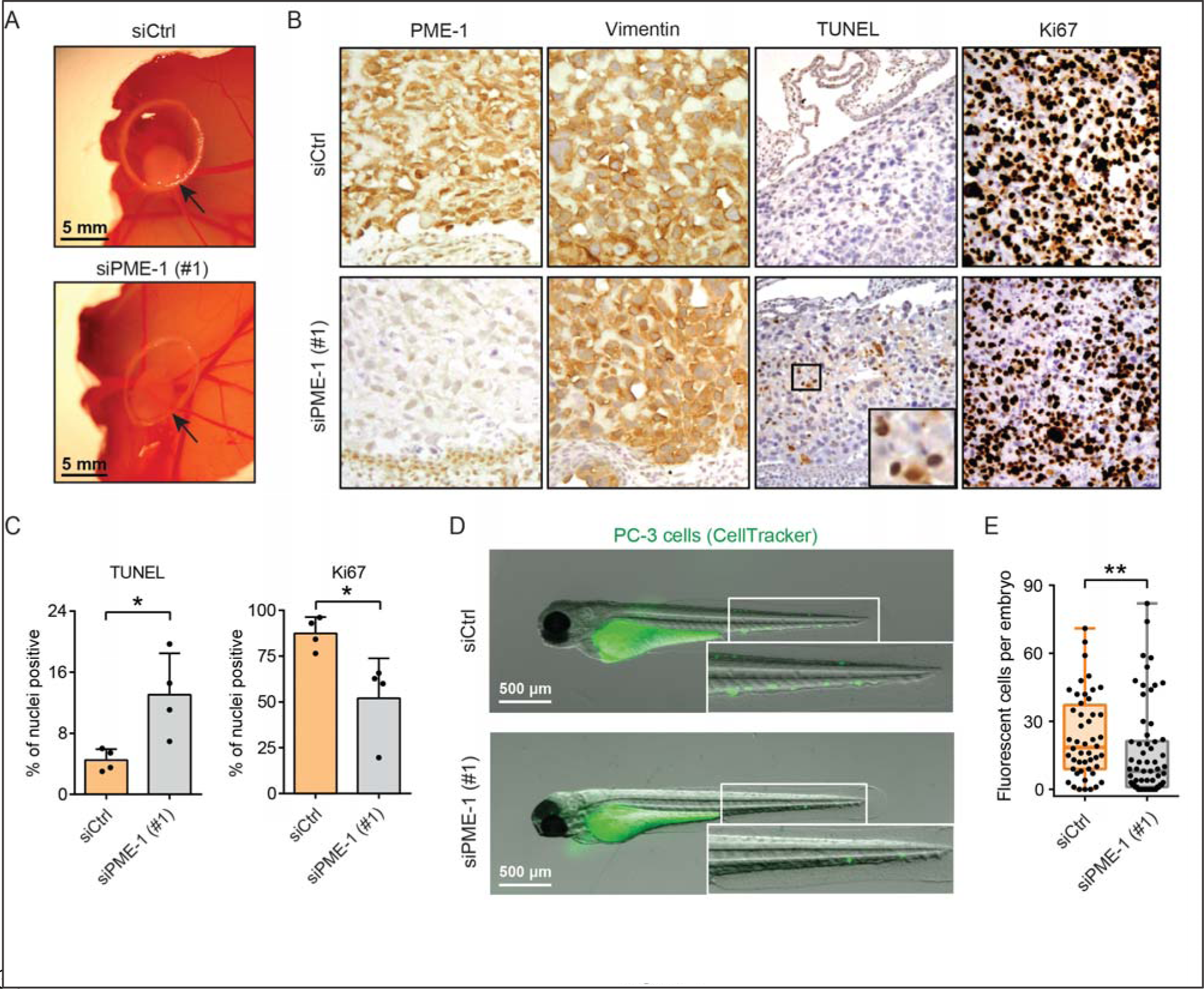
PME-1 supports *in vivo* anoikis resistance and survival of prostate cancer cells in circulation. **A**. The effect of PME-1 on anchorage-independent growth of PC-3 cells on chick chorioallantoic membrane (CAM). PC-3 cells were transiently transfected with either siCtrl or siPME-1, and 24 h post-transfection placed on the CAM. Growth of tumors was followed for 3-5 days. **B**. Immunohistological staining of dissected tumors using antibodies for PME-1, Vimentin, TUNEL and Ki67. **C**. Quantification of TUNEL- and Ki-67-positive nuclei in the excised CAM tumors. Mean ± SD, *p < 0.05, Mann-Whitney test. **D-E**. The survival of siCtrl- and siPME-1-transfected PC-3 cells in circulation. The cells were microinjected into the common cardinal veins of zebrafish embryos 72 h post-transfection. After overnight incubation the embryos were imaged by fluorescence stereomicroscopy (D). The number of surviving fluorescent tumor cells per embryo (E). Box plots depicting the range, 25th, 50th and 75th percentiles of the data, overlaid with the individual data points. **p < 0.01, Mann-Whitney test.

Anoikis resistance is crucial for tumor cell dissemination through circulation. To test whether high PME-1 expression could also suppress PCa cell death in circulation *in vivo*, we examined the survival of control and PME-1-depleted, fluorescent PC-3 cells microinjected into the common cardinal vein of zebrafish embryos [30]. Stereomicroscope imaging and quantification indicated that control cells were detected more frequently in the vasculature after 24 hours compared to the PME-1-silenced cells (Figure 4D, E), demonstrating that PME-1 supported survival of circulating PC-3 cells. These results are indicative of PME-1-mediated anoikis resistance of PCa cells *in vivo*.

### The potential role of histone H3 methylation in PME-1-mediated anoikis resistance

A recent phosphoproteome study identified 185 PME-1-regulated phosphopeptides in 128 proteins [31], indicating PME-1 control of multiple cellular pathways and regulatory programs in cancer cells. Thereby, the anoikis sensitivity of PME-1-depleted cells is expected to be governed by several different mechanisms. To identify any such putative target mechanism, we chose to interrogate the impact of PME-1 on mechanisms acknowledged to be involved in anoikis regulation [32, 33].

Related to the established role of focal adhesion kinase (FAK) signaling downstream of integrins in anoikis suppression [35], we performed an immunofluorescence analysis of phosphorylated FAK in control and PME-1-silenced cells that were grown on soft (0.5 kPa) hydrogels for 24 h. Even in low attachment conditions, endosomal FAK signaling can suppress anoikis and promote cell viability [17]. However, PME-1 silencing did not affect the overall p-FAK levels or existence of cytoplasmic p-FAK foci in these same culture conditions which resulted in anoikis sensitization in PC-3 cells (Figures 5A,B and 3D). Next, we analyzed the effects of PME-1 silencing on the phosphorylation of pro-survival kinase AKT and oncoprotein MYC by western blotting and by immunohistochemistry analysis of CAM tumors. However, no effects on either of these signaling mechanisms were observed upon PME-1 inhibition (Figure S3A,B). An additional candidate mechanism that could contribute to anoikis in PME-1-depleted cells was actin cytoskeleton remodeling upon cell rounding in low attachment conditions [32, 36]. To directly test whether disruption of the actin cytoskeleton would synergize with PME-1 inhibition to induce apoptosis, control (n.t. gRNA) and PME-1 KO (PME-1 gRNA) cells were treated with increasing concentrations of cytochalasin D that prevents actin polymerization (Figure 5B). However, western blotting analysis after 24 h of treatment with cytochalasin D did not show a PME-1-dependent difference in PARP cleavage, indicating that cell rounding and actin disruption are not causative mechanisms for inducing anoikis in PME-1-depleted cells (Figure 5C).

**Figure 5.**
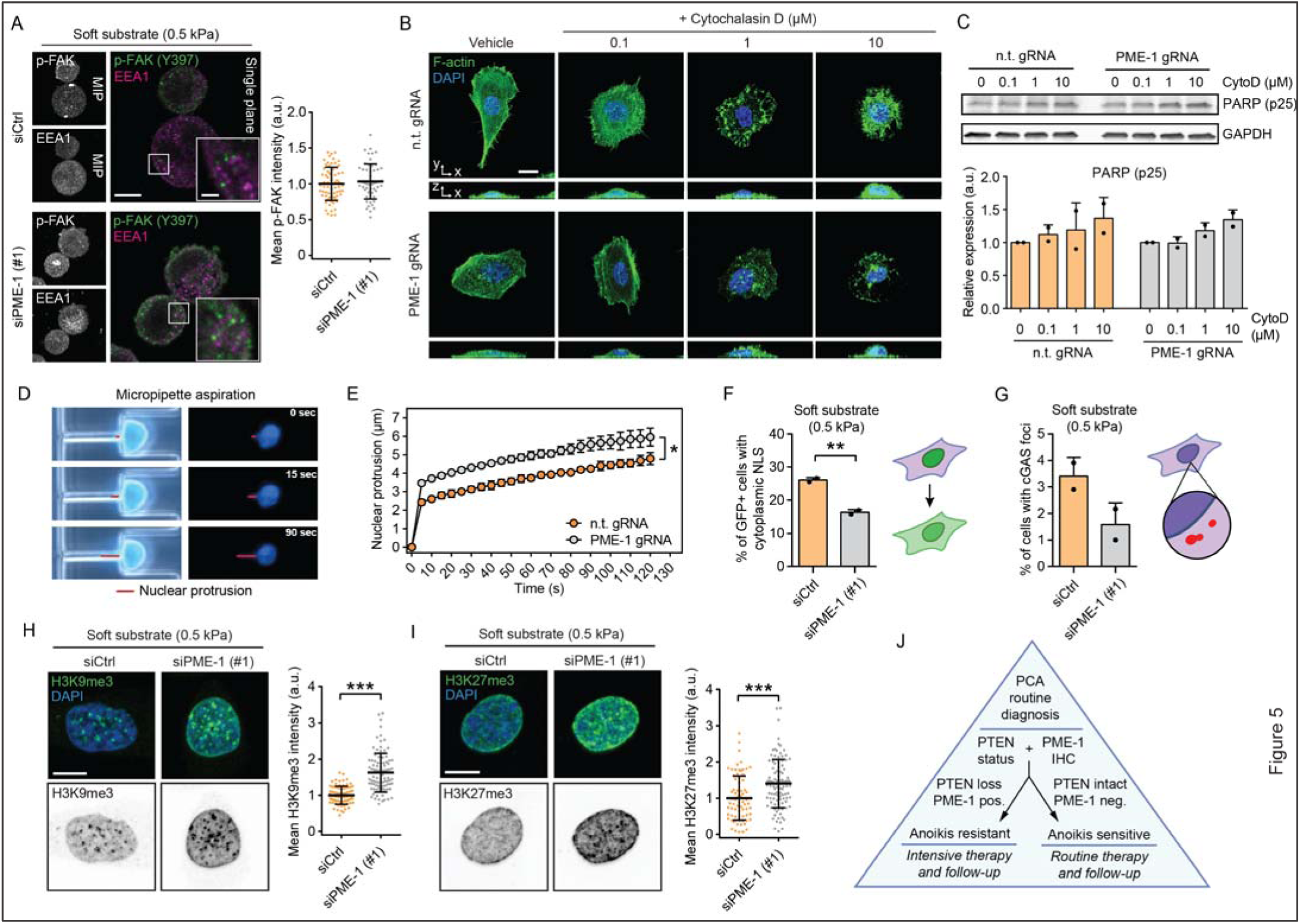
The potential role of histone H3 trimethylation in PME-1-mediated anoikis resistance. **A**. Immunofluorescence images (left) and quantification (right) depicting phosphorylated FAK (Y397) and early endosomal marker EEA1 in control and PME-1-depleted PC-3 cells grown on soft (0.5 kPa) hydrogels for 24 h. Scale bar, 10 µm (main), 2 µm (inset). Mean ± SD. **B.** Fluorescence images depicting filamentous actin in control and PME-1 KO PC-3 cells treated with DMSO or increasing concentrations of Cytochalasin D (0.1-10 μM). Scale bar, 20 µm. **C.** Western blot (top) and quantification (bottom) displaying PARP cleavage in PC-3 cells upon actin cytoskeleton disruption with Cytochalasin D. Mean ± SD of two independent experiments. **D-E.** Images (D) and quantification (E) showing nuclear deformation in PC-3 cells subjected to micropipette aspiration, as described in [34], over the course of two minutes. Mean ± SD of three independent experiments. *p < 0.05, Welch’s t-test. **F-G**. Quantification of GFP-NLS-positive cells with cytoplasmic GFP signal (F) and the percentage of transfected cells with visible cGAS-mCherry foci (G), respectively, in siCtrl- and siPME-1-treated cells grown on soft (0.5 kPa) substrate. Both readouts serve as a surrogate measure of compromised nuclear envelope. Mean ± SD of two independent experiments. **p < 0.01, Welch’s t-test **H-I**. Immunofluorescence images and quantification depicting H3K9me3 (H) and H3K27me3 (I) levels in transiently PME-1-depleted and control PC-3 cells, 48 h post-transfection and 24 h after seeding on soft (0.5 kPa) substrate. Scale bar, 10 µm. Mean ± SD, representative of two independent experiments. ***p < 0.001, Mann-Whitney test. **J**. Schematic representation of a putative model for including PME-1 and its role in anoikis sensitivity/resistance in PCa diagnostics and therapy decision.

To identify additional candidate mechanisms linked to PME-1-related anoikis regulation, we surveyed the most enriched cellular processes impacted by PME-1 depletion based on the phosphoproteome data [31]. Interestingly, nuclear envelope assembly and chromatin remodeling were the two most significantly PME-1-regulated cellular processes (Table S2). Notably, PME-1 physically associates with the nuclear lamina (NL) ([37] and Figure S4A,B) and regulates phosphorylation of lamins, key structural components of the NL, but also other related proteins such as LAP2A/B that connect the NL to chromatin (Figure S4C). It has previously been shown that NL directly influences chromatin architecture [38, 39], and that de-regulation of the NL can lead to DNA damage and cell death, limiting both cancer cell growth in soft agar and metastasis [38, 40, 41]. Therefore, we next assessed whether PME-1 inhibition could alter nuclear structure and stability. To this end, nuclear deformability of control and PME-1 KO cells were measured by subjecting them to a microfluidic micropipette aspiration experiment as previously described [34]. Notably, PME-1 KO cells had significantly more deformable nuclei, implying structural differences (Figure 5D,E). We then assessed whether these changes in nuclear mechanics would correlate with an increase in the number of harmful nuclear envelope ruptures [42] in PME-1-silenced cells. For this, we employed transient expression of GFP-tagged nuclear localization signal (NLS), and mCherry-tagged cyclic GMP–AMP synthase (cGAS), a cytosolic DNA sensor [42]. Surprisingly, PC-3 cells cultured in low stiffness conditions displayed fewer GFP-positive cytoplasms and less cytoplasmic cGAS foci, indicative of fewer compromised nuclear envelopes, upon PME- 1 silencing (Figure 5F,G and S5A-C). Together these results indicate that regardless of the newly identified role for PME-1 in regulating NL proteins (Figure S4C) and nuclear mechanics (Figure 5D-E), the anoikis sensitivity of PME-1-depleted PC-3 cells cannot be explained directly by apoptosis-inducing nuclear envelope disruptions.

However, PME-1-mediated alterations in different nuclear lamina proteins (Figure S4C) could also impact cell viability indirectly, by regulating chromatin structure and transcription (Table S2). Chromatin condensation is dependent on intact nuclear lamina, and heterochromatin structures are linked to the lamina via trimethylated histone H3 (H3K9me3 and H3K27me3) [39, 43, 44]. On the other hand, increased H3K27me3 and H3K9me3 levels sensitize cancer cells to apoptosis [21, 22, 45]. Thus, we studied whether PME-1 depletion would affect H3K9me3 and H3K27me3 levels in PC-3 cells grown on soft hydrogels. Indeed, PME-1 depletion was found to increase both H3K9me3 and H3K27me3 levels on soft substrate, concordant with the increased apoptosis observed in similar conditions (Figure 5H,I and 3D). Thereby, an increase in histone H3 methylation, and its subsequent apoptosis promoting effects [21, 22, 45], are likely contributing to the increased anoikis sensitivity in PME-1-depleted PCa cells. However, the direct mechanistic link between PME-1 and histone H3 methylation remains to be identified.

## DISCUSSION

Here, we discover a clinically relevant role for the PP2A inhibitory protein PME-1 in supporting anchorage-independent growth of PCa cells with compromised PTEN. Our data indicate that the anchorage-independence of PCa cells with concomitant inhibition of the two tumor suppressor phosphatases, PP2A and PTEN [3, 9], can largely be explained by their resistance to anoikis. Although anoikis suppression is a generally relevant mechanisms for tumor progression [14, 32], it may be of particular clinical importance in slowly progressing cancers such as PCa, where tumors can be diagnosed in the indolent phase, and there is a strong need to be able to both predict and inhibit the likelihood of disease progression (5).

Integrins act as key survival signaling receptors suppressing anoikis [18]. Here, we demonstrate that the onset of anoikis in PCa requires concomitant loss of both integrin-mediated matrix rigidity-dependent signaling and PME-1-mediated PP2A inhibition. Given the complexity of both signaling programs, the contributing molecular mechanisms are expected to be numerous, impeding the identification of a single key mechanism. Nevertheless, we could exclude several classical anoikis-related mechanisms from being involved in the increased anoikis sensitivity of PME-1-inhibited PCa cells with relative confidence. On the other hand, our results demonstrating a significant increase in apoptosis-promoting H3K9 and H3K27 trimethylation [21, 22, 45] do provide an interesting clue for the underlying mechanism. Although the direct mechanistic link between PME-1 and histone H3 methylation remains to be addressed in future studies, both the phoshoproteome data and increased nuclear deformability suggest that it could be attributed to abnormal NL-chromatin mechanics in the PME-1- deficient PCa cells.

PTEN and PP2A have both been identified independently as PCa tumor suppressor phosphatases (5, 12, 16, 52), but the clinical relevance of their co-operation has not been studied thus far. PTEN deficiency has been shown to promote anoikis resistance of PCa cells (20) and our data indicate that concomitant inhibition of a second tumor suppressor phosphatase PP2A by PME-1 renders the PTEN-deficient cells particularly well protected from anoikis. We hypothesize that this is likely to contribute to the observed clinical aggressiveness of these cancers. Mechanistically, PTEN-mediated anoikis resistance is mediated by AKT signaling (20-21), whereas we showed that PME- 1 depletion had no effect on AKT phosphorylation (Figure S3A,B). This indicates non-overlapping downstream mechanisms for PTEN and PME-1 in anoikis suppression, further explaining their synergetic actions. As PTEN genetic status can be routinely evaluated in current clinical PCa diagnostic practice (5), our results indicate a diagnostic utility of assessment of PME-1 status for patients with complete PTEN loss. Definition of PME-1 protein expression as a surrogate for PP2A activity status in tissues samples would greatly simplify biomarker analysis of PP2A function because it sidesteps the need for analysing all the possibly relevant subunits. Although further studies are clearly needed to validate these conclusions, our results indicate that patients with PTEN^loss^/PME-1^high^ tumors might benefit from more intensive follow-up, and/or from more aggressive therapies as first, and second line treatments (Figure 5J).

Together these results identify anoikis resistance as a candidate mechanism by which PME-1-mediated PP2A inhibition promotes malignant progression of PTEN-deficient PCa. Together with emerging orally bioavailable PP2A reactivating compounds with profound antitumor activity in *in vivo* PCa models [46], these results clearly emphasize the future importance of comprehensive understanding of PP2A biology for management of aggressive PCa. Future studies would also be needed to validate whether the functional co-operation of PTEN and PP2A extends to other cancers beyond PCa, and whether these findings would open novel therapeutic opportunities for simultaneously targeting both of these tumor suppressor phosphatases [47].

## ACKNOWLEGDEMENTS

We thank The Proteomics Core, The Cell Imaging and Cytometry Core, and Zebrafish Core at Turku Bioscience Centre supported by the University of Turku and Biocenter Finland. Taina Kalevo-Mattila and the rest of the Turku Bioscience personnel are acknowledged for their excellent technical help. This work was supported, in part, by grants from Academy of Finland (331237 to J.W and 325464 to J.I.), Academy of Finland CoE for Translational Cancer Research (J.I), Sigrid Juselius Foundation (J.W, J.I.), Finnish Cancer Institute (J.I.), the Finnish Cultural Foundation (A.I.), the Finnish Cancer Organization (J.W, J.I., P.T.), the National Institutes of Health [R01 GM137605, R01HL082792, and U54CA210184 to JL], the Department of Defense Breast Cancer Research Program [Breakthrough Award BC150580 to JL], and the National Science Foundation [CAREER Award CBET-1254846 and G14945R7301740 to JL]. This work was performed in part at the Cornell NanoScale Science & Technology Facility, a member of the National Nanotechnology Coordinated Infrastructure, which is supported by the National Science Foundation (Grant NNCI-2025233). A.I. has been supported by the University of Turku Doctoral Programme in Molecular Life Sciences (DPMLS).

## MATERIAL AND METHODS

### Cell lines

PC-3 and DU-145 prostate cancer cell line were obtained from ATCC and cultured in RPMI-1640. Pten^Δ/ Δ^; Trp53^Δ/ Δ^; lsl-tdTomato mouse embryonic fibroblasts were provided by Lloyd Trotmann and cultured in DMEM. All growth media were supplemented with 10% heat-inactivated FBS (Biowest), 2 mmol/L L-glutamine, penicillin (50 units/mL) and streptomycin (50 μg/mL). Cells were regularly tested and confirmed to be mycoplasma-negative, and were grown at 37 °C in a humidified atmosphere of 5% CO_2_.

### siRNAs

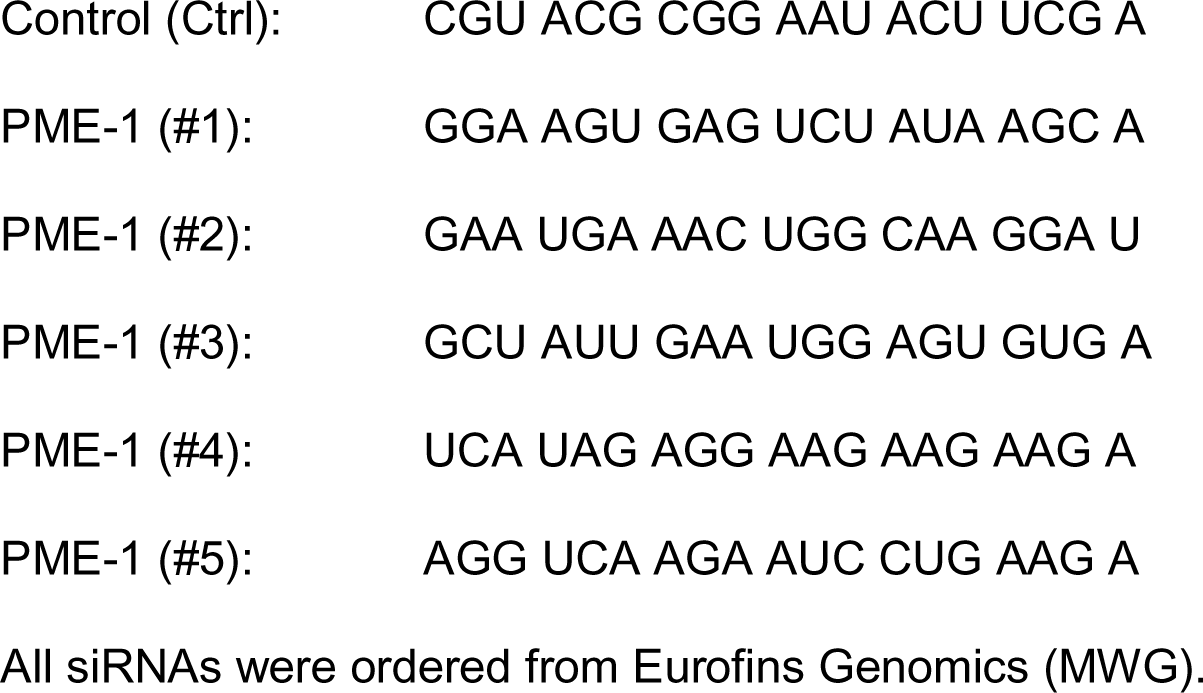

### Antibodies

The following antibodies were used at the indicated dilutions: PME-1, Santa Cruz Biotechnology sc-20086 (H-226), Western blotting 1:1000; PME-1, Santa Cruz Biotechnology sc-25278 (B-12), Immunohistochemistry 1:1000, Immunofluorescence 1:100; cleaved PARP-1, Abcam ab32064 [E51], Western blotting 1:1000; GAPDH, HyTest 5G4-6C5, Western blotting 1:5000; c-MYC, Abcam ab32072 [Y69], Western blotting 1:1000; p-Myc S62, Abcam ab78318, Western blotting 1:1000; Akt1/2/3, Cell Signalling Technology #9272, Western blotting 1:2000; p-Akt S473, Cell Signalling Technology #4060, Western blotting 1:1000; Lamin A/C, Santa Cruz Biotechnology sc- 7292 (636), Immunofluorescence 1:250; EEA1, Santa Cruz Biotechnology sc-137130 [G4], Immunofluorescence 1:100; p-FAK Y397, Cell Signalling Technology #8556, Immunofluorescence 1:100; Histone H3K9me3, Cell Signalling Technology #13969 [D4W1U], Immunofluorescence 1:500; Histone H3K27me3, Cell Signalling Technology #9733 [C36B11], Immunofluorescence 1:500.

### siRNA/plasmid transfection

For siRNA transfections, Lipofectamine RNAiMAX (Thermo Fisher Scientific) was used following the manufacturer’s instructions. 2.5×10^5^ cells were seeded on a 6-well plate one day before transfection to reach 60-70% confluency. Cells were then transfected with 25 pmol siRNA and 7.5µl RNAiMAX per well and assayed 48-72h after transfection, unless otherwise stated. For plasmid transfections, Lipofectamine 3000 (Thermo Fisher Scientific) was used following the manufacturer’s instructions. 3-4×10^5^ cells were seeded on a 6-well plate one day before transfection to reach 80-90% confluency. Cells were then transfected with 2.5 µg DNA, 5 µl P3000 reagent and 3.75 µl Lipofectamine 3000 per well and assayed 24-48h after transfection, unless otherwise stated.

### Colony formation assay

Cells were reseeded at low confluency (1-4×10^3^) on a 12-well plate 24 h after siRNA transfection and grown for around 10 days to allow the formation of cell colonies. Colonies were then fixed with ice-cold methanol and stained with 0.2% crystal violet solution (in 10% ethanol) for 10 minutes at room temperature. Excess stain was removed by repeated washing with PBS. Plates were dried and scanned with Epson perfection V700 scanner. Quantifications were performed with ColonyArea ImageJ plugin [48], and graphs were plotted using the area % values.

### Anchorage-independent colony formation assay

For the anchorage-independent colony formation assay, which typically correlates with *in vivo* tumorigenicity, 2×10^4^ cells were resuspended in 1.5 ml of growth medium containing 0.4% agarose (4% Agarose Gel, Gibco; top layer) and plated on 1 ml bottom layer containing growth medium and 1.2% agarose in a 12-well plate. After 14 days of growth, colonies were stained overnight with 1 mg/ml Nitro blue tetrazolium chloride (NBT; Molecular Probes) in PBS. Colonies were imaged using a Zeiss SteREO Lumar V12 stereomicroscope. Analysis was done using the ImageJ software. First, the background was subtracted using the rolling ball function with a radius of 50 μm, then auto-thresholding was applied to separate the colonies. Area percentage was calculated using the ImageJ built-in function ‘Analyze Particles’. Particles smaller than 500 μm^2^ were not considered colonies, and were excluded from the analysis.

### Western blotting

Western blot protein lysates were prepared in 1x RIPA buffer (150 mM sodium chloride, 1.0% NP-40, 0.5% sodium deoxycholate, 0.1% SDS and 50 mM Tris, pH 7.5) containing PhosSTOP™ phosphatase inhibitors and cOmplete™ EDTA-free protease inhibitors (Roche). DNA in the protein samples was sheared by sonication and the amount of protein was estimated using Pierce™ BCA Protein Assay Kit (Thermo Fisher Scientific). Lysates were separated on 4–20% Mini-PROTEAN® TGX™ Gels (Bio-Rad) and transferred by wet blotting to PVDF membranes (Millipore), or via semi-dry transfer to nitrocellulose membranes (Bio-Rad). Unspecific antibody binding was blocked with 5% non-fat dry milk in TBST. Incubation for the primary antibodies was performed overnight at 4 °C in either 5% non-fat dry milk, or in 5% BSA for the phospho-specific antibodies. For detection, HRP-labeled secondary antibodies (DAKO) followed by incubation with Pierce™ ECL Western Blotting Substrate (Thermo Fisher Scientific) were used. Alternatively, LI-COR Biosciences secondary antibodies (IRDye 680 or IRDye 800) were used, followed by detection by Odyssey® Imaging Systems or Bio-Rad Laboratories ChemiDoc Imaging Systems.

### Generation of Pten^Δ/Δ^; Trp53^Δ/Δ^ mouse embryonic fibroblasts

Pten^Δ/Δ^; Trp53^Δ/Δ^; lsl-tdTomato MEFs were generated as previously described (56). To stably overexpress PME-1 in these cells, PME-1 cDNA was cloned into pCSCIW-2 lentiviral construct which was then packaged into lentiviral particles. Following transduction, GFP positive cells were sorted and overexpression of PME-1 was confirmed by western blotting.

### CRISPR/Cas9-mediated PME-1 knock-out

To generate PME-1 knock-out cells, PC-3 cells were transduced with lentivirus containing lentiCas9-Blast construct (Addgene #52962) and selected with growth medium containing 4 µg/ml blasticidin for approximately one week. Afterwards, the cells were infected with lentivirus sgRNA against PME-1 exon 3 (sequence: ACTTTTCGAGTCTACAAGAGTGG) cloned into pKLV-flipedU6gRNA_PB_BbsI_PGKpuro2ABFP construct. After puromycin selection (2 µg/ml in growth medium), PME-1 KO on the protein level was confirmed by western blotting. Single cell clones were obtained by cell sorting and the knock-out was confirmed by sequencing (Primers. Forward CACCGCTTTTCGAGTCTACAAGAGGT; reverse TAAAACGAAGATCTGTCTGCAGAAAC). Cas9- and non-targeting gRNA-expressing cells served as a negative control in the functional assays.

### Chick chorioallantoic membrane (CAM) assay

To start chick embryonic development for the chorioallantoic membrane assay, fertilized eggs were kept rotating in an incubator at 37 °C and 50-60% humidity for four days. After the initial incubation, a small hole was introduced at the sharp edge of the egg and sealed with parafilm. Following four more days of incubation on stationary racks, the hole was enlarged and a plastic ring was placed on top of blood vessels in the chorioallantoic membrane. Next, 1×10^6^ PC-3 cells in 20 µl volume of a 1:1 mixture of ice-cold PBS and matrigel were pipetted inside the ring on the membrane. The hole was then covered with parafilm and the eggs were incubated for three more days. At day twelve of embryonic development, the animals were sacrificed by freezing the eggs for 15 min and the tumor cell mass was dissected from the membrane and processed for further analysis.

### Immunohistochemistry

Hematoxylin/eosin staining and immunohistochemistry were performed on 3-μm-thick sections of 4% paraformaldehyde-fixed and paraffin-embedded tissues. Following rehydration, endogenous peroxidase was blocked by incubation in 50% MetOH, 1% H_2_O_2_. Subsequent antigen retrieval was performed with the 2100 Retriever (Aptum) in R-Universal Buffer. Unspecific antibody binding was blocked with 10% goat serum in 2% BSA/PBS prior to overnight incubation at 4°C with the primary antibody. For detection, the DAKO EnVision peroxidase system, followed by incubation with 0.01% diaminobenzidine (Sigma) was used.

### Zebrafish *in vivo* dissemination assay

The xenotransplantation of zebrafish embryos was performed as described in detail in [30] with some modifications. PC-3 cells were washed with PBS, stained with CellTracker Green CMFDA dye (5 µM, Thermo Fisher Scientific) and detacher and detached with trypsin-EDTA in a single incubation step at 37 °C. Subsequently, cells were pelleted by centrifugation and washed with PBS twice. This was followed by filtration through 40 µM mesh into FACS tube (BD Falcon, 352235) and pelleting cells by centrifugation. Finally, cells were resuspended into 30 µl of injection buffer (2% PVP in PBS) and kept on ice until transplanted. Zebrafish (*Danio rerio*) of pigmentless casper strain (*roy^-/-^ ; mitfa^-/-^*) [49] was used in the experiments under licence no. MMM/465/712-93 (issued by Finnish Ministry of Forestry and Agriculture) and following legislation: the European Convention for the Protection of Vertebrate Animals used for Experimental and other Scientific Purposes and the Statutes 1076/85 and 62/2006 of The Animal Protection Law in Finland and EU Directive 86/609. The embryos were obtained through natural spawning, and were cultured in E3-medium (5 mM NaCl, 0.17 mM KCl, 0.33 mM CaCl_2_, 0.33 mM MgSO_4_) at 33°C. At 2 or 3 days post-fertilization, the embryos were anesthesized with 200 mg/ml Tricaine and embedded in 0.7% low-melting point agarose. Subsequently, the cell suspension was microinjected into common cardinal vein (duct of Cuvier) of the embryo using glass capillaries (Transfertip), CellTramVario microinjector and InjectMan micromanipulator (all from Eppendorf). Embryos were liberated from the agarose gel using forceps and successfully transplanted embryos were selected to the experiment. After overnight incubation at 33 °C, the embryos were anesthesized again with Tricaine and imaged using Zeiss StereoLumar V12 fluorescence stereomicroscope. The number of surviving cells was counted manually from the images using ImageJ.

### Proximity ligation assay (PLA)

Cells were grown to approximately 80% confluency on sterilized coverslips, fixed for 10 min in 4% PFA and permeabilized for 10 min with 0.5% Triton X-100 in TBS. The subsequent steps were completed following the manufacturer’s instructions (Sigma). Briefly, unspecific antibody binding was blocked with the provided blocking solution for 30 min. The slides were then incubated with the primary antibodies overnight at 4 °C, with the mouse and rabbit probes for one hour at 37 °C, with the ligation mix for 30 min at 37 °C and the amplification mix for 100 min at 37 °C. The washing steps in-between the individual steps were carried out with Buffer A (Sigma) and the final washing step with Buffer B (Sigma). Slides were mounted in Mowiol mounting medium and imaged with a Zeiss LSM780 confocal microscope.

### Investigation of nuclear envelope integrity

PC-3 cells were transfected with PME-1 or control siRNA as indicated above. After 24 h, the same cells were transfected with both pCDH-CMV-NLS-copGFP-EF1-blastiS and pCDH-CMV-cGASE225A/D227A-mCherry2-EF1-blastiS [42] in a 1:1 ratio and incubated overnight. Finally, the cells were detached, transferred onto 0.5 kPa hydrogels and grown for an additional 24 h before fixing. Lamin-A,C, NLS-GFP, cGAS-mCherry and DNA were visualized using (immune)fluorescence microscopy. After thresholding, individual cells were stratified and quantified based on the presence/absence of cytoplasmic NLS and cGAS foci.

### Cell-matrix interaction assay

In order to quantitatively control the degree of cell-matrix interaction, 1 mg/ml poly-L-lysine-grafted polyethylene glycol (PLL(20)-g[3.5]-PEG(2), from SuSoS AG) and 1 mg/ml 50% biotinylated PLL-g-PEG (PLL(20)-g[3.5]-PEG(2)/PEG(3.4)-biotin(50%), from SuSoS AG) stock solutions were mixed with 10 mM Hepes (pH 7.4) to create 0.1 mg/ml working solutions with 0, 0.1, 0.25, 1, 5 or 20% biotinylated PLL-g-PEG. A plastic 96-well plate was coated with each of the different stock solutions for 1 h at room temperature and washed twice with PBS. Next, the coated wells were immersed in approximately 250 ng/cm^2^ of streptavidin-conjugated (Abnova FastLink Streptavidin Labeling Kit) fibronectin fragment, FNIII(7-10), in PBS and incubated for 1 h at room temperature. Afterwards, the wells were washed three times with PBS. In order to evaluate the degree of cell adhesion to the coated wells, PC-3 cells were detached using HyQTase (HyClone, SV30030.01) and 1×10^4^ cells were seeded into each well. The cells were left to adhere and spread for 30 min, after which the wells were washed twice with PBS while avoiding draining them completely. The cells were fixed with 4% PFA/PBS for 5 min, washed three times with PBS and visualized by incubating the wells in 0.2% (w/v) crystal violet (Sigma)/10% ethanol for 15 min at room temperature. The wells were washed three times with PBS, air dried, and 100 µl of 10% acetic acid per well was used to resolubilize the stain. The results (A595) were read using a spectrophotometer (Multiskan Ascent, Thermo Fisher Scientific).

In order to investigate the degree of anoikis in PC-3 cells grown on surfaces with varying amounts of integrin ligand, cells that had been transfected with control or PME-1- targeting siRNAs approximately 48 h prior to the experiment were seeded on a new 96- well plate coated with 0-20% biotinylated PLL-g-PEG and streptavidin-FNIII(7-10), at 5×10^3^ cells per well. Each well was supplemented with 1:200 Annexin V-FITC (eBioscience), and additional wells with 20% biotinylated PLL-g-PEG and FNIII(7-10) were supplemented with 1 µM staurosporine (Merck Millipore) to serve as a positive control for apoptosis. The plate was imaged every hour for >2.5 days and analyzed for the presence of Annexin V-positive cells and debris using IncuCyte S3 (Essen Bioscience).

### Actin cytoskeleton disruption

Plastic 6-well plates and 8-well microscopy slides (Ibidi µ-Slide) were coated overnight at 4 °C with 2.5 µg/ml of bovine plasma fibronectin (Merck-Millipore, 341631) and collagen type I (Sigma, C8919). On the following day, control and PME-1 KO PC-3 cells were seeded on the 6-well plates and microscopy slides at approximately 30% seeding density and supplemented with DMSO (control) or 0.1 10 µM of Cytochalasin D (Sigma, C8273-1MG) three hours later. Cells on the microscopy slides were fixed after two-hour treatment and processed into fluorescence microscopy samples as indicated below. The remaining cells were lysed after 24 hours and analyzed for the presence of cleaved PARP by western blotting.

### Immunofluorescence

Cells were fixed with 4% PFA for 10 min at room temperature, and simultaneously permeabilized and blocked with 0.3% Triton in 10% horse serum (Gibco) for 15 min at room temperature. All samples were incubated in primary antibodies (diluted in 10% horse serum) overnight at 4 °C, and stained with secondary antibodies for 1-2 h at room temperature on the following day. Appropriate Alexa Fluor-conjugated secondary antibodies (Thermo Fisher Scientific) were used at a 1:300 dilution in PBS. Nuclei were counterstained with DAPI (4’,6-diamidino-2-phenylindole) and filamentous actin with Alexa Fluor 488-conjugated phalloidin (Thermo Fisher Scientific).

### Polyacrylamide hydrogels

35 mm glass bottom dishes (MatTek Corporation, P35G-1.0-14-C) were treated with 100 µl of Bind-silane solution [7.14% Bind-silane (Sigma, M6514) and 7.14% acetic acid in absolute ethanol] for 15 min, washed twice with absolute ethanol and left to dry completely. A pre-polymer mix comprising 5.4% acrylamide (Sigma) and 0.043% N,N -′ methylenebisacrylamide (Sigma) in PBS was prepared to obtain hydrogels with an elastic modulus of appoximately 0.5 kPa [50] Polymerization was initiated by adding 2.5µl of 20% ammonium persulfate (Bio-Rad) and 1 µl of N,N,N ′,N′ - tetramethylethylenediamine (Sigma). The solution was vortexed, 13 µl was added on top of the glass bottom dish, a 13 mm glass coverslip was placed on the drop and the gel was left to polymerize for 1 h at room temperature. After polymerization, the dish was filled with PBS and the coverslip was carefully removed. Hydrogels were made permissive for protein binding by incubating them in 500 µl of Sulfo-SANPAH solution [0.2 mg/ml Sulfo-SANPAH (Sigma, 803332) and 2 mg/ml N-(3-dimethylaminopropyl)-N′- ethylcarbodiimide hydrochloride (Sigma, 03450) in 50 mM HEPES] for 30 min on slow agitation, followed by a 10 min UV exposure (∼30 mW/cm^2^, 253.7 nm). Activated gels were washed three times with PBS to get rid of residual Sulfo-SANPAH. Alternatively, pre-activated polyacrylamide hydrogels of variable stiffness were ordered from Matrigen Life Technologies. 35 mm glass bottom dishes (SV3510-EC-0.5) were used for imaging and 6-well plates (SW6-EC-0.5, SW6-EC-50) for growing cells for lysis. All hydrogels were functionalized with bovine plasma fibronectin and collagen type I by incubating the dishes in 5 µg/ml of each protein for 1-2 h at 37 °C, or overnight at 4 °C, before use.For western blot analysis of cells on hydrogels, 0.5 and 50 kPa hydrogel-coated 6-well plates were immersed in cell culture medium for 30 min and seeded with 300,000 PME- 1 or control siRNA-treated PC-3 cells per well. The cells were grown on the gels for 24 h. Once the cultures reached 80-90% confluency, the cells were washed once and scraped into cold PBS, spun down and lysed in RIPA buffer as described above. For immunofluorescence experiments, PC-3 cells were seeded on 0.5 kPa hydrogel-coated glass bottom dishes and grown for an additional 24 h before fixing and sample preparation.

### Microscopy and image analysis

Fluorescent specimen were imaged using a spinning disk or laser scanning confocal microscope. The spinning disk system used was a Marianas spinning disk confocal microscope with a Yokogawa CSU-W1 scanning unit, controlled by SlideBook 6 software (Intelligent Imaging Innovations). The objective used was a 40x/1.1 W LD C-Apochromat objective (Zeiss), and images were acquired using an Orca Flash4 sCMOS camera (Hamamatsu Photonics). The laser scanning confocal microscope used was an LSM780, controlled by Zen 2010 (Zeiss), and the objective used was a 40x/1.2 W C-Apochromat objective (Zeiss). Images were analyzed using ImageJ (National Institutes of Health) and CellProfiler (Broad Institute) softwares.

### Statistical analysis

Statistical analyses and plotting were performed using GraphPad Prism v6.05 (GraphPad). The names and/or numbers of individual statistical tests, samples and data points are indicated in figure legends. Unless otherwise noted, all results are representative of a minimum of two independent experiments and two-tailed p-values are reported.

