## Supplementary material for "PME-1 suppresses anoikis and is associated with therapy relapse of PTEN-deficient prostate cancers": Table 1

| Table 1. Univariate Analysis of PME IHC and clinical variables. | | | |
| --- | --- | --- | --- |
|  | PME IHC Low (n = 238) | PME IHC High (n = 82) | P-Value |
| Age at RP, years (mean, SD) (*n* = 320) |  |  |  |
| < 60 | 81 (34.0) | 24 (29.3) | 0.515 |
| 60 - 70 | 132 (55.5) | 46 (56.1) |  |
| > 70 | 25 (10.5) | 12 (14.6) |  |
| Preoperative PSA, ng/ml (n, %) (*n* = 254) |  |  |  |
| ≤10.0 | 100 (52.1) | 32 (51.6) | 0.995 |
| 10.1-20.0 | 59 (30.7) | 19 (30.6) |  |
| >20.0 | 33 (17.2) | 11 (17.8) |  |
| Grade group at RP (n, %) (*n* = 320) |  |  |  |
| 1 | 74 (31.1) | 5 (6.1) | **< 0.001** |
| 2 | 65 (27.3) | 17 (20.7) |  |
| 3 | 73 (30.7) | 36 (43.9) |  |
| 4 | 22 (9.2) | 17 (20.7) |  |
| 5 | 4 (1.7) | 7 (8.5) |  |
| pT (n, %) (*n* = 299) |  |  |  |
| 2 | 150 (67.3) | 28 (36.8) | **< 0.001** |
| 3-4 | 73 (32.7) | 48 (63.2) |  |
| Lymph node status (n, %) (*n* = 314) |  |  |  |
| Negative | 228 (97.4) | 78 (97.5) | 1.000 |
| Positive | 6 (2.6) | 2 (2.5) |  |
| Secondary Therapy (n, %) (n = 320) |  |  |  |
| No | 171 (71.8) | 39 (47.6) | **< 0.001** |
| Yes | 67 (28.2) | 43 (52.4) |  |
| Death from any cause (n, %) (*n* = 320) |  |  |  |
| Yes | 104 (43.7) | 46 (56.1) | 0.055 |
| No | 134 (56.3) | 36 (43.9) |  |
| Death from prostate cancer (n, %) (*n* = 320) |  |  |  |
| Death due to PC | 20 (8.4) | 11 (13.4) | 0.197 |
| Alive, or dead from other causes | 218 (91.6) | 71 (86.6) |  |
| Pearson's chi-square | | | |
