## Supplemental figures 1-5 for "PME-1 suppresses anoikis and is associated with therapy relapse of PTEN-deficient prostate cancers"

A

PME-1 vs PTEN Status

| | PME-1 Low | PME-1 High | Total | P |
|---|---|---|---|---|
| PTEN Intact | 193 (81.1) | 57 (69.5) | 250 | **0.043**[a] |
| Complete PTEN Loss | 45 (18.9) | 25 (30.5) | 70 | |
| Total | 238 | 82 | 320 | |

[a] Fisher's exact test

PME-1 vs AR Status

| | PME-1 Low | PME-1 High | Total | P |
|---|---|---|---|---|
| AR Low | 97 (40.8) | 14 (17.1) | 111 | **< 0.001**[a] |
| AR High | 141 (59.2) | 68 (82.9) | 209 | |
| Total | 238 | 82 | 320 | |

PME-1 vs ERG Status

| | PME-1 Low | PME-1 High | Total | P |
|---|---|---|---|---|
| ERG Negative | 128 (53.8) | 27 (32.9) | 155 | **0.001**[a] |
| ERG Positive | 110 (46.2) | 55 (67.1) | 165 | |
| Total | 238 | 82 | 320 | |

B

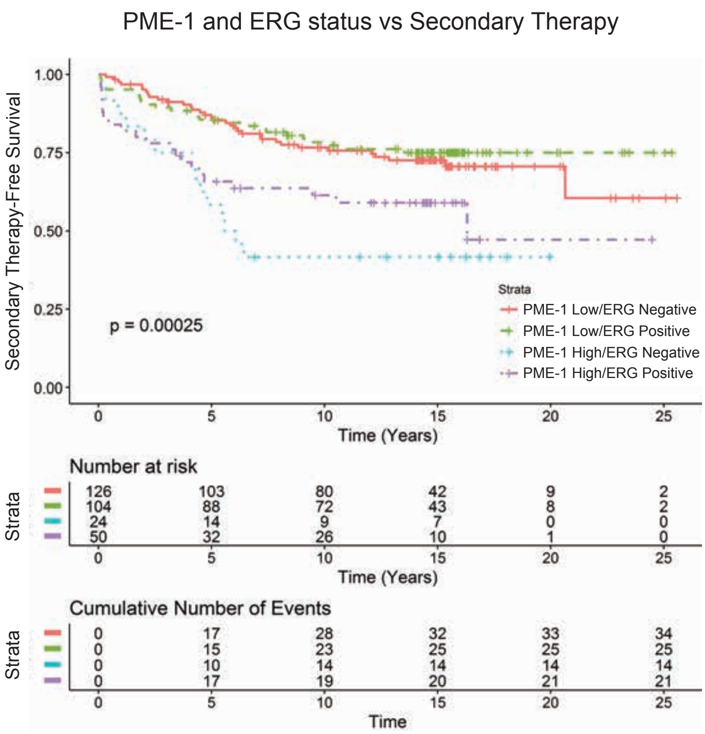


Figure S1

Figure S1. High PME-1 expression associates with total PTEN loss in prostate cancer patient samples A. PME-1 status was correlated to previously assessed PTEN, AR and ERG status. PME-1 expression significantly associate with complete PTEN loss, but also with high AR expression and ERG positivity status, as analysed by Fisher's exact test. B. Kaplan-Meyer analysis of time to secondary therapies after primary treatment, based on PME-1 status in combination with ERG.

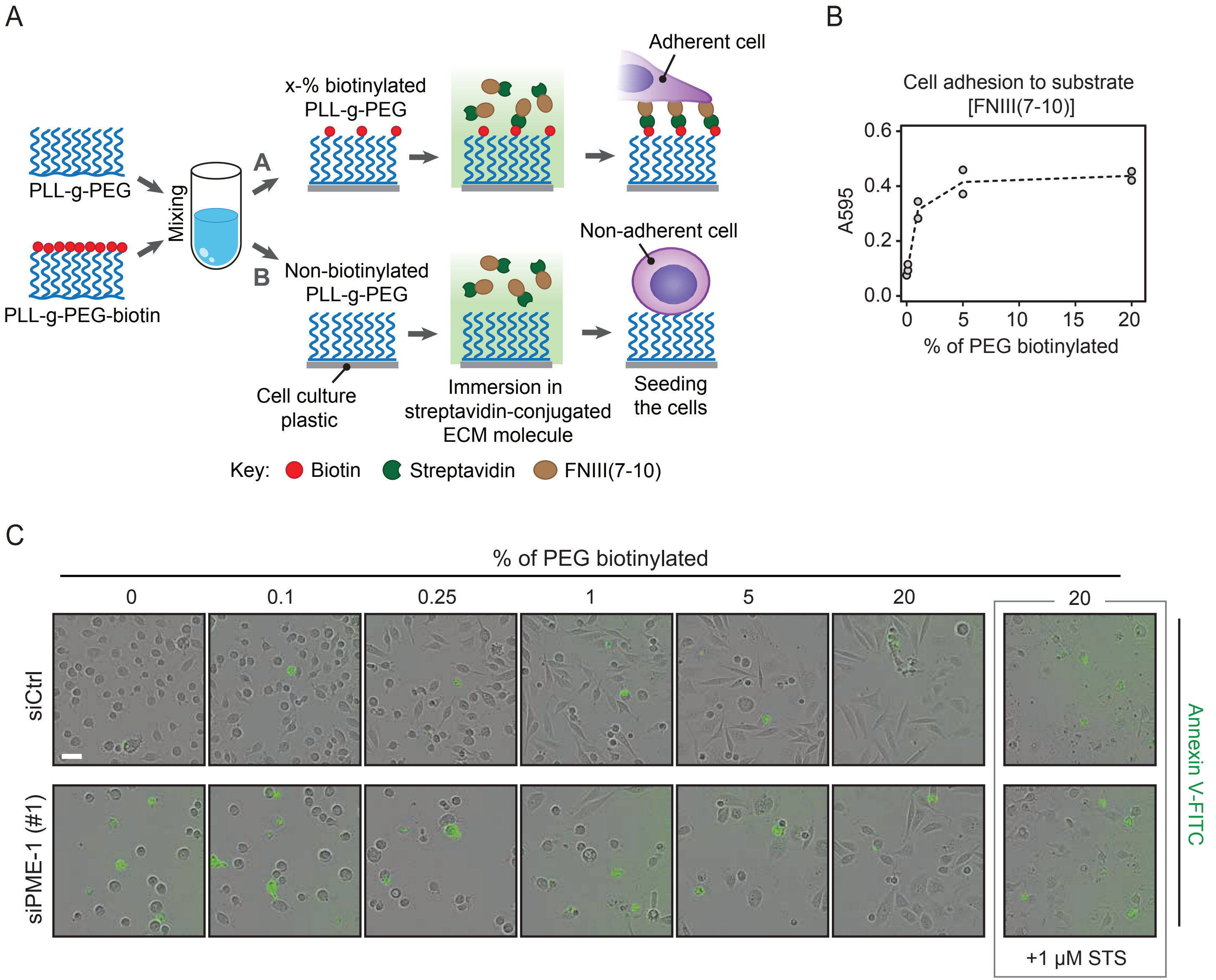


Figure S2

Figure S2. Modulation of PC-3-integrin ligand interaction using biotinylated PLL-g-PEG and streptavidin-conjugated fibronectin fragment.
A. Schematic representation of the preparation of PLL-g-PEG-coated tissue culture surfaces. Stock solutions of PLL-g-PEG and PLL-g-PEG-biotin were mixed together in 10 mM Hepes buffer (pH 7.4) to yield PLL-g-PEG-solutions with varying amounts of biotinylated compound (A), or no biotin groups at all (B). Next, biotinylated PLL-g-PEG was coupled to streptavidin-conjugated FNIII(7-10) to enable selective adhesion of cells to specific coated surfaces. B. Standard curve depicting PC-3 adhesion to PLL-g-PEG- and FNIII(7-10)-coated plastic over 30 min. Cell-substrate adhesion increases as a function of the fraction of biotinylated PEG. C. Phase contrast images and overlaid fluorescence data showing control and PME-1-depleted PC-3 cells, as well as Annexin V positive cells and debris after 60 h on PLL-g-PEG- and FNIII(7-10)-coated plastic. STS, staurosporine. Scale bar, 50 μm.

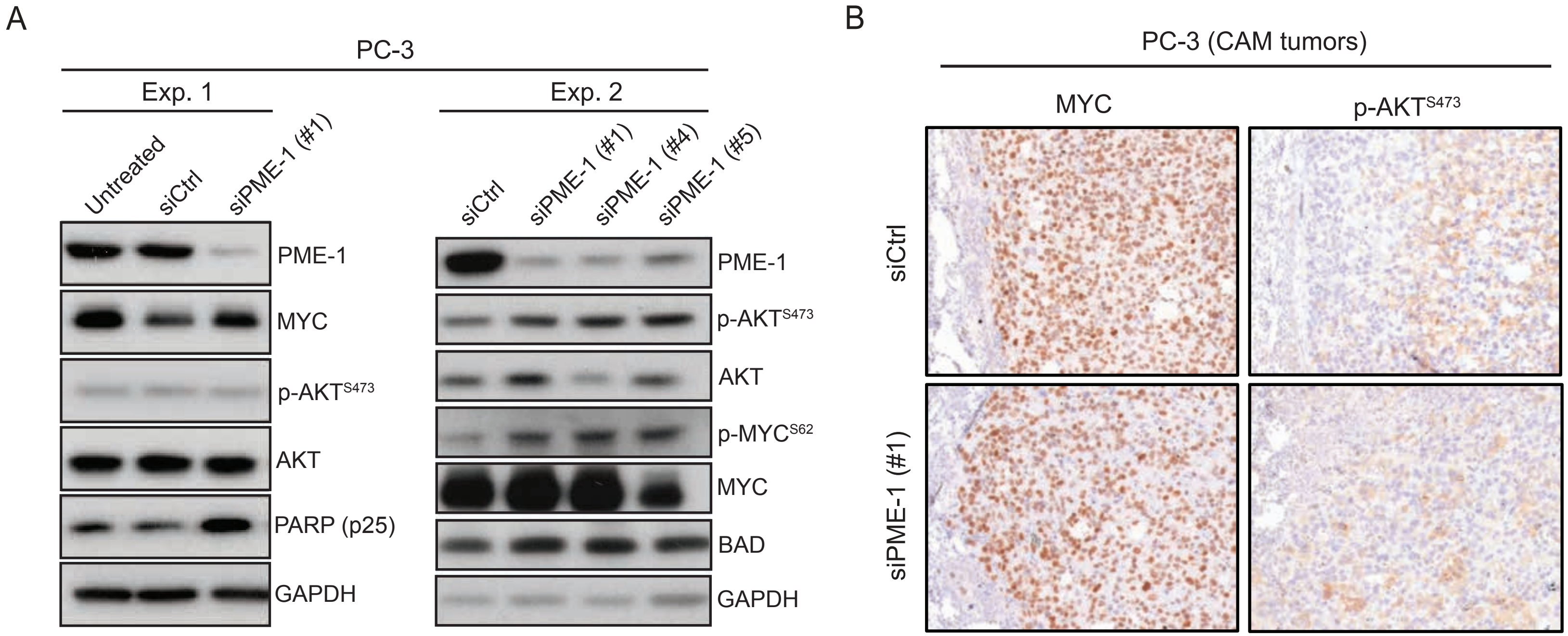


Figure S3

Figure S3. PME-1 is not affecting AKT or MYC signaling in PC-3 cells
A. The effect of PME-1 depletion upon transient transfections on AKT and MYC were analysed by western blotting.
B. The effect of PME-1 silencing on MYC levels and AKT phosphorylation was assayed by immunohistochemistry of CAM tumors.

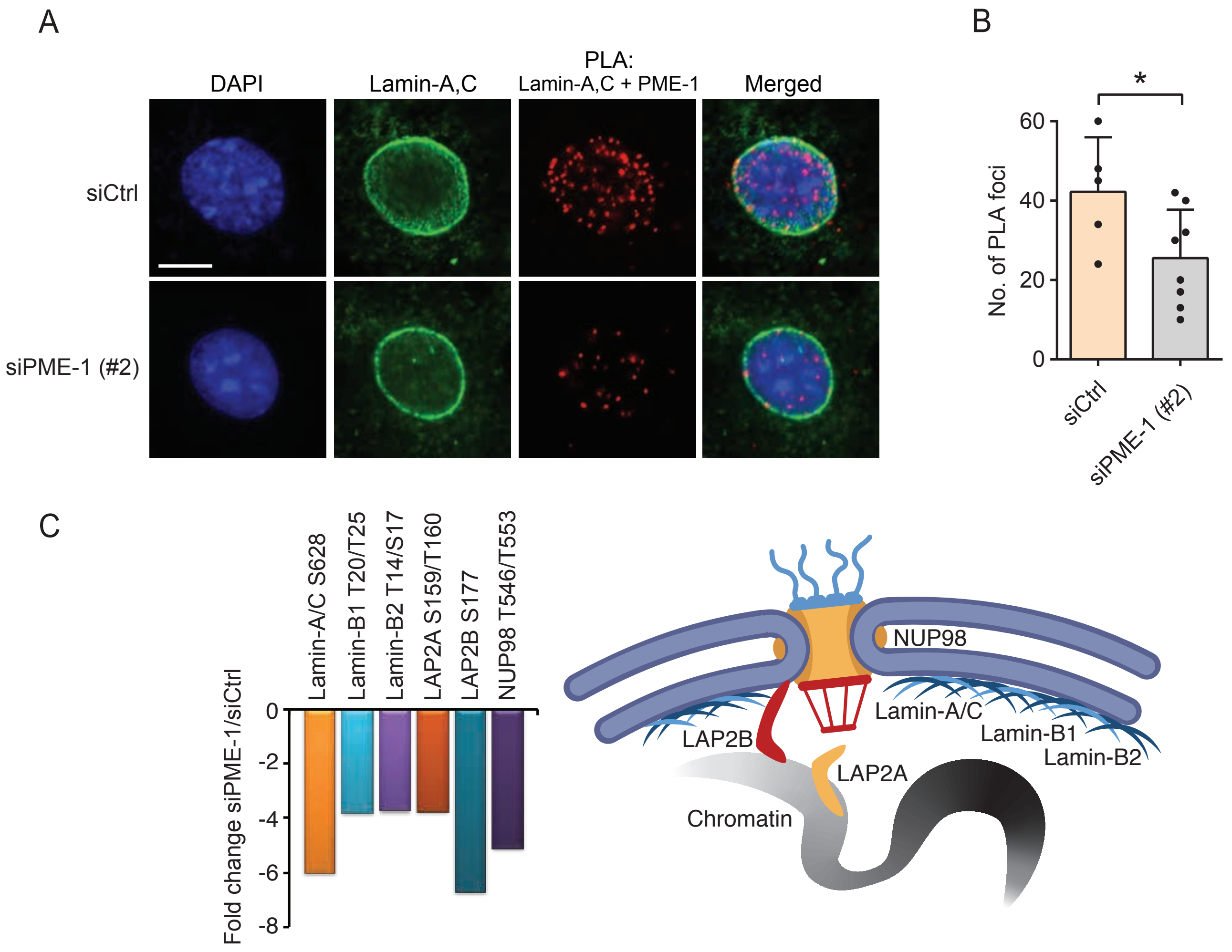


Figure S4

Figure S4. PME-1 co-localizes with Lamin-A/C and regulates the phosphorylation of multiple nuclear lamina components.
A. Proximity ligation assay (PLA) and immunofluorescence illustrate the co-localization of PME-1 with Lamin-A/C in siRNA-treated PC-3 cells. DNA was counterstained using DAPI. Scale bar, 10 µm. B. The number of PLA foci in siCtrl- and siPME-1-transfected cells. Mean ± SD, *$p < 0.05$, Mann-Whitney test. C. PME-1 was found to regulate the phosphorylation of lamins, LAP2A/B and NUP98 (36).

A

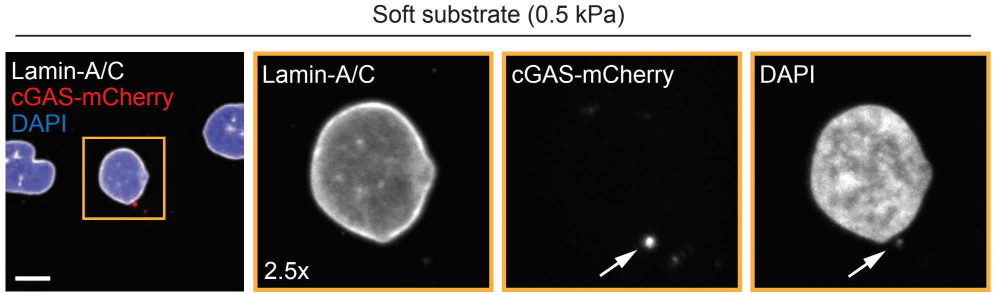


B

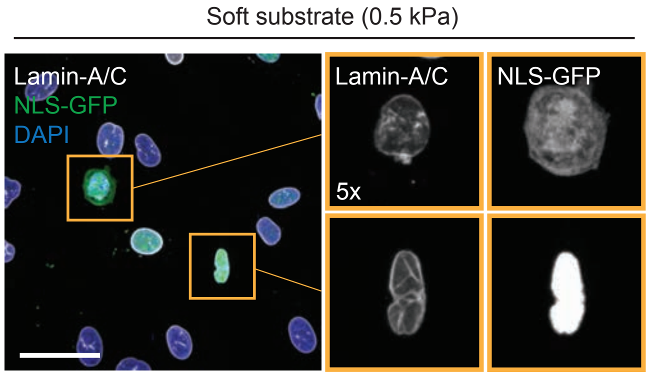


C

| siRNA | NLS-GFP cytoplasmic (%) | | p-value (Chi-squared) |
|---|---|---|---|
| | – | + | |
| Ctrl | 152 (74.1) | 53 (25.9) | |
| PME-1 | 155 (83.8) | 30 (16.2) | **0.020** |

| siRNA | cGAS-mCherry foci (%) | | p-value (Chi-squared) |
|---|---|---|---|
| | – | + | |
| Ctrl | 1131 (96.7) | 38 (3.3) | |
| PME-1 | 1009 (98.5) | 15 (1.5) | **0.007** |

Figure S5

Figure S5. PME-1 silencing does not compromise PC-3 nuclear envelope integrity on soft substrates.
A. Immunofluorescence image depicting Lamin-A/C, cytoplasmic DNA and a corresponding cGAS-mCherry aggregate (white arrows) in a siCtrl-transfected PC-3 cell on soft (0.5 kPa) hydrogel. Scale bar, 10 µm. B. Immunofluorescence image showing Lamin-A/C and NLS-GFP in siCtrl-transfected PC-3 cells on soft (0.5 kPa) hydrogel. ROI: some of the cells present with cytoplasmic NLS-GFP (top), implying a compromised nuclear envelope, while in others the protein is constrained solely in the nucleus (bottom). Scale bar, 50 µm. C. Contingency tables depicting all the GFP positive cells analyzed for NLS localization (top) and all the transfected cells analyzed for the presence of cGAS-mCherry foci (bottom).
Pooled results from two independent experiments.
