## Supplementary material for "PME-1 suppresses anoikis and is associated with therapy relapse of PTEN-deficient prostate cancers": Table S1

| Table S1. Demographics of the radical prostatectomy patient cohort. | |
| --- | --- |
|  | Total Cohort (n=358) |
| Age at RP, years (median, IQR) (*n* = 358) | 64.0 (59.2 – 67.9) |
| Preoperative PSA, ng/ml (n, %) (*n* = 283) |  |
| ≤10.0 | 143 (50.5) |
| 10.1-20.0 | 89 (31.4) |
| >20.0 | 51 (18.0) |
| Grade group at RP (n, %) (*n* = 358) |  |
| 1 | 93 (26.0) |
| 2 | 93 (26.0) |
| 3 | 114 (31.8) |
| 4 | 45 (12.6) |
| 5 | 13 (3.6) |
| pT (n, %) (*n* = 334) |  |
| 2 | 202 (60.5) |
| ≥3 | 132 (39.5) |
| Lymph node status (n, %) (*n* = 352) |  |
| Negative | 342 (97.2) |
| Positive | 10 (2.8) |
| Follow-up time after RP, years (median, range) (n=358) | 15.7 (0.7-28.6) |
| Death from any cause (n, %) (*n* = 358) | 172 (48.0) |
| Death from prostate cancer (n, %) (*n* = 358) | 33 (9.2) |
| Patients receiving secondary therapy after RP (n, %) (*n* = 358) | 124 (34.6) |
